# Mitochondrial introgression suggests extensive ancestral hybridization events among *Saccharomyces* species

**DOI:** 10.1101/028324

**Authors:** David Peris, Armando Arias, Sandi Orlić, Carmela Belloch, Laura Pérez-Través, Amparo Querol, Eladio Barrio

## Abstract

Horizontal Gene Transfer (HGT) in eukaryotic plastids and mitochondrial genomes is common, and plays an important role in organism evolution. In yeasts, recent mitochondrial HGT has been suggested between *S. cerevisiae* and *S. paradoxus*. However, few strains have been explored due to the lack of accurate mitochondrial genome annotations. Mitochondrial genome sequences are important to understand how frequent these introgressions occur and their role in cytonuclear incompatibilities and fitness. In fact, most of the Bateson-Dobzhansky-Muller genetic incompatibilities described in yeasts are driven by cytonuclear incompatibilities. In this study, we have explored the mitochondrial inheritance of several worldwide distributed *Saccharomyces* species isolated from different sources and geographic origins. We demonstrated the existence of several recombination points in the mitochondrial region *COX2-ORF1*, likely mediated by the transfer of two different types of *ORF1* (F-*Sce*III), encoding a freestanding homing endonuclease, or mostly facilitated by A+T tandem repeats and regions of integration of GC clusters. These introgressions were shown to occur at intra- as well as at interspecific levels. This suggest a complex model of *Saccharomyces* evolution which involve several ancestral hybridization events in wild environments.

## 2. Introduction

Chloroplast and mitochondrial genomes are prone to introgressions, recombinations and Horizontal Gene Transfers (HGT) (Keeling 2009; Hao *et al*. 2010), which likely play an important role in the evolution of eukaryotes (Andersson 2009). Introgressions, recombination, and HGT are considered reticulate events due to their non-verticality character which violates the phylogenetic tree-like assumption (Nakhleh 2011). Instead, reticulate events are better represented by phylogenetic networks where a taxonomic group can be connected to multiple lineages contributing to it (Bapteste *et al*. 2013). Mitochondria is involved in multiple cellular processes (Hatefi 1985; Green and Reed 1998; Starkov 2008). In yeasts, around 750 nuclear encoded proteins must coordinate with those encoded in the mitochondrial genome (Sickmann *et al*. 2003). Indeed, a new interdisciplinary field, the “mitonuclear ecology”, is devoted to the study of the evolutionary consequences driven by mitonuclear conflicts (Hill 2015).

In yeasts, reticulate evolution in mitochondrial genomes, such as recombination, has been mostly focused on *Saccharomyces cerevisiae* (Dujon *et al*. 1974; Birky *et al*. 1982; Taylor 1986; MacAlpine *et al*. 1998), the model yeast species in the *Saccharomyces* genus (Hittinger 2013). The mechanism of mitochondrial genome inheritance is well known in *Saccharomyces* (Ling *et al*. 2007; Basse 2010; Ling *et al*. 2011); but despite the recently mitochondrial genome characterization of a hundred *S. cerevisiae*, mostly clinical (Wolters *et al*. 2015), and few *S. paradoxus*, and the detection of mitochondrial introgression between those two species (Wu *et al*. 2015; Wu and Hao 2015), little is known about the mitochondrial genome inheritance of most *Saccharomyces* species and their hybrids.

In this study, we inferred the mitochondrial inheritance of globally distributed 517 *Saccharomyces* strains and 49 natural interspecific hybrids using the *COX2* mitochondrial gene marker. This gene has been successfully used either for phylogenetic purposes (Belloch *et al*. 2000; Kurtzman and Robnett 2003) and for the identification of mtDNA inheritance (Peris *et al*. 2012a, 2012b, 2014, 2016; Badotti *et al*. 2013; Pérez-Través *et al*. 2014; Rodríguez *et al*. 2014). The *COX2* sequence region was prone to contain highly incongruent data which indicates reticulate evolution. To understand this incongruence, we extended the study by sequencing the *COX2* neighbor *ORF1* locus, from 72 representative strains. The *ORF1* gene is potentially encoding a putative free-standing homing endonuclease. Most authors consider *ORF1* is not a truly functional gene and they do not annotate it in the vast majority of recent *Saccharomyces* mitochondrial genome annotations. A homing endonuclease gene (HEG) is a selfish element described to evolve neutrally, following three steps: i) invasion of an empty site (HEG^−^) (Colleaux *et al*. 1986; Burt and Koufopanou 2004); ii) the disruption of the selfish gene by the continuous accumulation of mutations, which generate premature stop codons, or by the invasion of GC clusters, which introduces frameshifts that can disrupt their coding frame; iii) and the final loss of the gene (Goddard and Burt 1999). *ORF1* was classified as a putative homing endonucleases from the LAGLIDADG family, due to the presence of the typical P1 and P2 LAGLIDADG domains (Bordonné *et al*. 1988). In this protein family, the homing endonucleases are required to maintain the LAGLIDADG domains. These domains play an important role during protein folding and the catalytic activity (Belfort and Roberts 1997). The incongruence found at the end of *COX2* gene might be linked to the life cycle of the *ORF1*.

To better define mitochondrial inheritance, we sequence a better mitochondrial marker, *COX3*. The *COX3* gene, which encodes for cytochrome c oxidase subunit III, was shown as a better marker for describing the mitochondrial inheritance of the vast majority of the seventy-two representatives of the *Saccharomyces* lineages, selected based on *COX2* haplotypes. This gene supported the recombinations found in the *COX2-ORF1* region. All these data showed a complex model of *Saccharomyces* evolution in wild and industrial environments.

## 3. Material and Methods

### 3.1 Saccharomyces strains and culture media

A collection of worldwide distributed 517 *Saccharomyces* strains, and 49 natural hybrids (Figure 1, Table S1) were used in this study. Species assignment for each strain was delimited by different authors using molecular techniques, such as the sequencing of 5.8S-ITS region, Random Fragment Length Polymorphisms (RFLP), a multilocus sequence approach or whole genome sequencing (Table S1). Hybrids were mostly characterized on the basis of restriction analysis of 35 nuclear genes (Table S1). *Naumovozyma castellii* sequences were included as the outgroup references to root phylogenetic trees. Yeast strains were grown at 28 °C in YPD (2% glucose, 2% peptone and 1% yeast extract).

**Figure 1.**
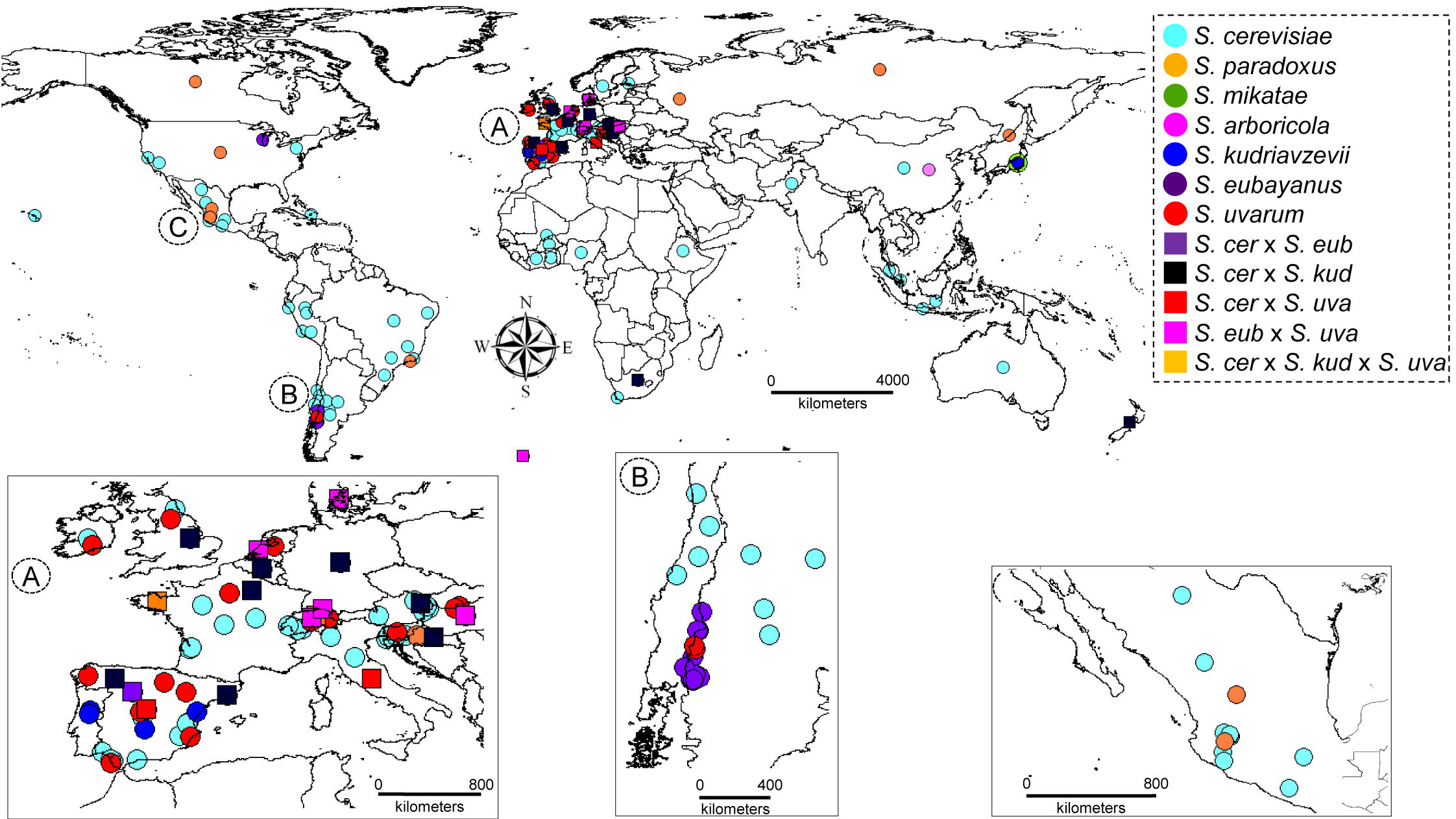
*Saccharomyces* worldwide distribution map. *Saccharomyces* strains were colored according to the species designation or the hybrid character (Table S1). Highly sampled regions were zoom in for a better resolution.

### 3.2 PCR amplification, sequencing and gene alignments

Total yeast DNA was extracted following the procedure described by Querol *et al*. (1992). 517 partial gene sequences (585bp) of the *COX2* mitochondrial gene were amplified by PCR using the primers described by Belloch *et al*. (2000). *COX3* and *ORF1* gene sequences were amplified and sanger-sequenced using primers described in Table S2, or retrieved from public databases as indicated below for a total of seventy-two *Saccharomyces* strains, which represented the most frequent *COX2* haplotypes (Table S1). The *ORF1* sequencing followed a primer walking approach. *ORF1* gene amplification from IFO1815 failed to be amplified, and the *ORF1* sequence from the reference *S. eubayanus* strain (FM1318) was not available in the period of this study; however, a recent study demonstrated that *S. eubayanus ORF1* from this strain is in advance stay of decay (Baker *et al*. 2015). Sequences were deposited in GenBank under Accession nos. JN676363-JN676823 and JN709044-JN709115. Gene sequence accession numbers from previously sequenced strains are shown in Table S1. Sequences from *S. cerevisiae* strains from the *Saccharomyces* Genome Resequencing Project v1 (SGRP v1) were retrieve using the blast server (http://www.sanger.ac.uk/cgibin/blast/submitblast/s_cerevisiae_sgrp).

*COX2* and *COX3* sequences were aligned using `CLUSTALW`, as implemented in `MEGA v5` (Tamura *et al*. 2011), and manually trimmed. For *ORF1* we used `MUSCLE` (Edgar 2004) to align the aminoacid sequences and the nucleotide sequence alignment was further refined by visual inspection in `Jalview 4.0b2` (Waterhouse *et al*. 2009). Annotations and polymorphism figures were done in `Geneious v. R6` (Kearse *et al*. 2012).

### 3.3 *COX2, ORF1* and *COX3* haplotype classification, and *COX*2 genetic diversity

`DnaSP v5` (Librado and Rozas 2009) was used to calculate the number of haplotypes of *COX2, ORF1* and *COX3*, and genetic statistics of *COX2*, such as the number of polymorphic sites(s), average number of differences between sequences (k), nucleotide diversity (π) and haplotype diversity (Hd) based on the species designation.

### 3.4 Phylogenetic analysis and detection of recombination

*COX2, ORF1* and *COX3* phylogenetic tree reconstructions were performed in `MEGA v5` (Tamura *et al*. 2011) by using the Neighbor-Joining (NJ) method, with 10000 pseudoreplicates bootstrapping for branch support. *COX2* and *COX3* median joining networks were reconstructed using `PopART v1.7.2b` (http://popart.otago.ac.nz). *COX2* and *ORF1* Neighbor-Net phylogenetic networks were also reconstructed using `SplitsTree v4` (Huson and Bryant 2006) to explore the presence of incongruence in our dataset.

An alignment with representative sequences of each *COX2* haplotype was used as an input for `RDPv3.44` (Martin *et al*. 2010) to detect and define recombination points. We reported significant recombination points (p-value < 0.05) detected by two or more methods implemented in `RDPv3.44`, after applying a Bonferroni correction for multiple comparisons. Although, different recombination points were detected, we defined two *COX2* segments using the most frequent recombinant sites. *COX2* gene was divided into two segments referred as 5’-end (positions 1-496 in the alignment [124-620 in the reference *COX2* gene sequence from the S288c strain, SGD ID: S000007281]) and 3’-end (from 527-end of alignment [651-708 in the reference *COX2* sequence]). The Maximum Likelihood (ML) phylogenetic trees for both *COX2* segments were reconstructed with the best fitted models, inferred using `jModeltest` (Posada 2008). `Tree Puzzle v5.2` (Schmidt *et al*. 2002) was used to test the phylogenetic congruence of the two inferred phylogenetic trees to the species *Saccharomyces* phylogenetic tree topology (Borneman and Pretorius 2015). The statistical significance of these comparisons was performed by the Shimodaira-Hasegawa (Shimodaira and Hasegawa 1999) and ELW (Expected-Likelihood Weights) (Strimmer and Rambaut 2002) tests.

A concatenated alignment of *COX2* position 621 to the end and the partial *ORF1* sequences was generated. Indels, mostly due to AT repetitive regions, and GC clusters were removed, generating a dataset of ~1.9 Kb, used for further analyses. For the detection of recombinant sites, we followed the approach described above. Four segments were described for the most frequent recombinant points. First segment (224bp) takes the *COX2* region and the 246 nucleotide positions of *ORF1* (corresponding to nucleotide 292 in the S288c *ORF1* gene, SGD ID: S000007282). The last nineteen nucleotides of *COX2* are the first *ORF1* nucleotides due to an overlap of both coding DNA sequences (CDS) (Bordonné *et al*. 1988). The second segment goes from position 247 to 644 of the *ORF1* (from 203 to 704 in S288c *ORF1)*, the third was built with the region covering from 645 to 920 nucleotide positions (706-980 in S288c), and the fourth segment goes from 921 to the end of the alignment (981-1435 in S288c). Recombinant segments were also supported by the `GARD` (Genetic Algorithm Recombination Detection) method (Kosakovsky Pond *et al*. 2006b) implemented in `Datamonkey` (Delport *et al*. 2010). Basically, `GARD` method seeks for the most supported number of partitions to avoid that recombinant regions affect further phylogenetic analysis. In a second step, `GARD` delimits the recombinant points and checks that phylogenetic tree topologies, which are reconstructed using each significant partition, are statistically different by performing a Shimodaira-Hasegawa test (Shimodaira and Hasegawa 1999; Kosakovsky *et al*. 2006a).

A new concatenated alignment (~2.3 Kb) containing *COX2, ORF1*, and *COX3* was generated. Using `RDP` and `GARD` we reanalyzed the dataset for exploring congruence with the previous analysis or the detection of new recombinant regions.

### 3.5 Species tree reconstruction

The *Saccharomyces* species tree was reconstructed by using a concatenated alignment (~1.2 Kb) generated with `FASCONCAT` v1.0 (Kück and Meusemann 2010), consisting of *RIP1* (511 bp), *MET2* (469 bp) and *FUN14* (294 bp) gene sequences. The strains included into the species tree were selected as representatives of each lineage of their corresponding population, including the type strains for *S. paradoxus* (CBS432), *S. mikatae* (IFO1815), *S. kudriavzevii* (IFO1802), *S. arboricola* (CBS10644), *S. uvarum* (CBS7001), and *S. eubayanus* (FM1318, monosporic derivative of CRUB1568). The type strain of *S. cerevisiae* S288c was not included due to its known mosaic genome (Liti *et al*. 2009). The genes were selected based on the sequence availability for all representative of each population/species analyzed in this study. Gene sequences were retrieved from Genbank, SGRP, using the online blast database for *S. arboricola* (http://www.moseslab.csb.utoronto.ca/sarb/blast/), or local `BLAST` databases (Altschul *et al*. 1997) generated with the Saccharomyces Sensu Stricto (SSS) genomes (http://www.saccharomycessensustricto.org/cgi-bin/s3.cgi?data=Assemblies&version=current). A ML phylogenetic tree was inferred using `RAxML v8` (Stamatakis 2014) by performing 100 heuristic searches for the best gene tree, which branch support was approximated by bootstrapping 1000 pseudosamples.

### 3.6 Statistical analysis

X^2^ test for detecting country or isolation source bias distributions for *S. cerevisiae* strains in the *COX2* haplogroups was performed in R statistical package (Adler and Murdoch D 2009). X^2^ test was also applied to detect a significant association between the recombinant sites and recombinogenic nucleotide sequences (GC clusters, A+T tandem repeats, and the starting *ORF1* CDS).

## 4. Results

### 4.1 Incongruence data in *COX2* gene sequence suggests reticulate events among *Saccharomyces* species

To understand the mitochondrial inheritance of worldwide distributed wild and domesticated *Saccharomyces* yeasts (Figure 1), we sequenced and retrieved from public databases the mitochondrial *COX2* gene sequences, generating a set of 566 *Saccharomyces COX2* sequences (Table S1). *COX2* sequence alignment contained 80 phylogenetically informative positions (see details in Supplementary Text). A Median-Joining network showed thirteen haplogroups, supported by sequence inspection (Figure 2 and S1). *S. cerevisiae* were found in three haplogroups: C1a, C1b and C2. Three haplogroups differentiated *S. paradoxus* populations: P1 (Europe – CECT10308), P2 (America – 120MX) and P3 (Far East). Two haplogroups for *S. mikatae:* M1 (IFO1815) and M2 (IFO1816), *S. kudriavzevii* strains were split into K1 (European – ZP591, and Asia A – IFO1802) and K2 (IFO1803). Finally, *S. arboricola* (CBS10644) was contained in haplogroup *A, S. eubayanus* strains from Patagonia B, and A populations in haplogroup E, and Holarctic and South American *S. uvarum* strains in haplogroup U.

**Figure 2.**
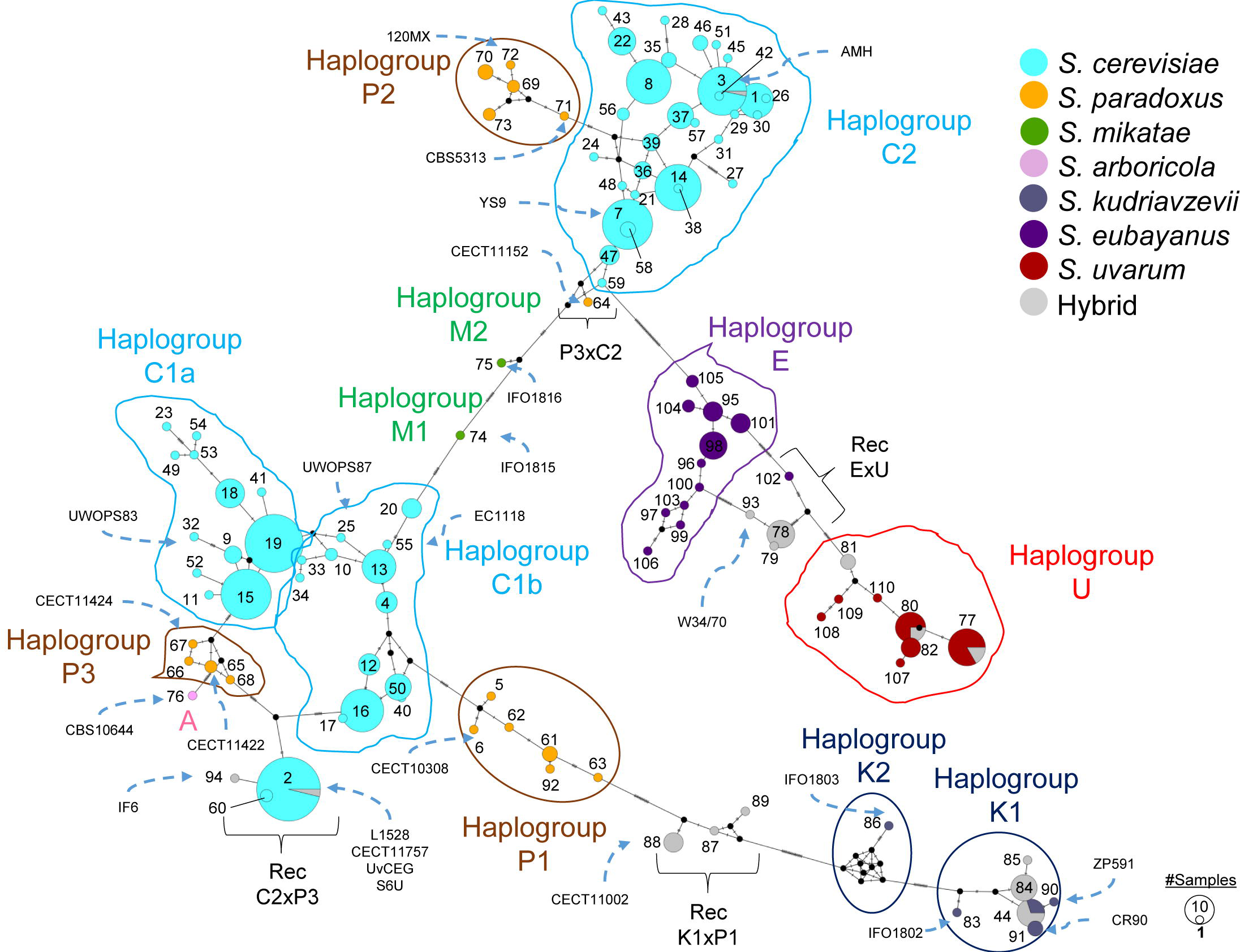
*COX2* Median-Joining phylogenetic network. A hundred and ten *COX2* haplotypes (Table S1) were represented by circles. Circle size is scaled according to the haplotype frequency. Pie charts show the frequency of each species in that particular haplotype. Number of mutations from one haplotype to another are indicated by lines in the edges connecting haplotypes. Strain names discussed in the main text are represented and connected to their corresponding *COX2* haplotype by a dashed blue arrow. Diversity statistics corresponding to the *COX2* sequence are found in Table S3.

The *COX2* phylogenetic tree failed to reconstruct the species tree (Figure 3) as expected from the observation of the MJ network (Figure 2). Analysis of recombination demonstrated the presence of several recombinant haplotypes among strains from different species (Figure S1). Clear recombinations were identified in Far Eastern *S. paradoxus* (haplotype 64, strain CECT11152, syn. IFO1804) and *S. cerevisiae* strains and hybrids (haplotypes 2, 60 and 94, which representatives are *S. cerevisiae* L1528, and CECT11757) (Figure 2 and S1). We segmented the original *COX2* alignment based on the most common recombination point detected by `RDP` (Figure S1), after excluding a region with the presence of homoplasic mutations. A Neighbor-Net (NN) phylogenetic network was reconstructed for each *COX2* segment (Figure S2). The phylogenetic network reconstructed using the 5’ end segment differentiated haplotypes by the species designation, except for American *S. paradoxus* (haplotypes 69, 70 and *71)*, which share identical sequences with some *S. cerevisiae* strains (Figure S2A). Haplotypes from natural hybrids (haplotypes 78, 79, 87, 88, 89 and 93) and *S. eubayanus* (haplotype 102) still appear in ambiguous positions in the network due to the presence of different recombination points than that selected as the common recombinant point (Figure S1). In any case, the *COX2* 3’ end segment (Figure S2B) was responsible of most of the incongruence.

**Figure 3.**
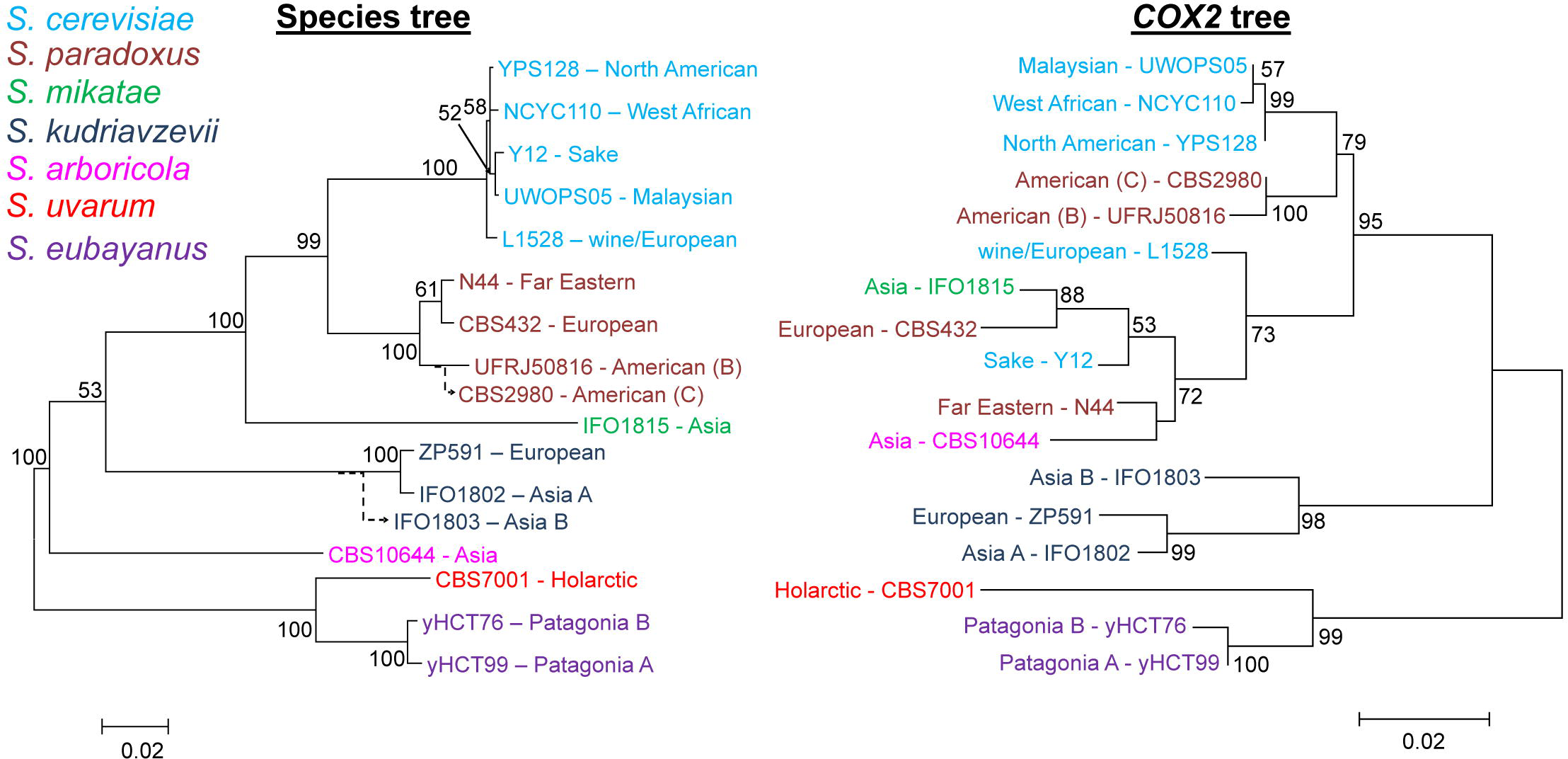
Species and *COX2* ML phylogenetic trees. ML phylogenetic tree using a concatenated alignment of *RIP1, MET2* and *FUN14* (left), and *COX2* (right) are represented. Strains were colored according to the species designation. CBS2980 and IFO1803 strain positions in the phylogenetic species tree were inferred according to Leducq *et al*. (2014) and Hittinger *et al*. (2010). Despite the small region (~1.2 Kb) used for representing the species tree, this topology is congruent with previous phylogenetic representations for individual species (Liti *et al*. 2009, Leducq *et al*. 2014, Hittinger *et al*. 2010, Peris *et al*. 2016), or studies including multiple species (Scannell *et al*. 2011, Borneman and Pretorius 2015, Boynton and Greig 2014). Bootstrap values above 50 are given for each branch. Scales are given in nucleotide substitution per site.

A careful inspection detected the potential donors of recombinant *COX2* sequences from hybrids. In the case of haplotypes corresponding to double, *S. cerevisiae* × *S. kudriavzevii*, which representative is CECT11002, and a triple hybrid *S. cerevisiae* × *S. kudriavzevii* × *S. uvarum* (haplotypes 87, 88, and 89), one of the donors was identified as *S. kudriavzevii* (European or Asian), and surprisingly the second donor was a European *S. paradoxus* (Figure S1). Hybrids *S. cerevisiae* × *S. kudriavzevii* isolated from wine (UvCEG) and a dietary supplement (IF6), and *S. cerevisiae* × *S. uvarum* isolated from wine (S6U) inherited an already recombinant mitochondrial genome from *S. cerevisiae* closely related to the wine strains *S. cerevisiae* L1528, and CECT11757 (Figure 2, S1, Table S1). Hybrids *S. eubayanus* × *S. uvarum*, and *S. cerevisiae* × *S. eubayanus* (haplotypes 78, 79, and 93, the latter including the well-known hybrid lager strain, W34/70) were also recombinant between *S. eubayanus* and *S. uvarum* as demonstrated previously by Peris *et al*. (2014).

*S. mikatae* IFO1815, and *S. kudriavzevii* IFO1803 haplotypes were also potentially recombinant (Figure S1). In the case of IFO1803, the amino acid sequence indicates a recombination between *S. kudriavzevii* and *S. uvarum* (Figure S1); although, the region donated by *S. uvarum* corresponds to an unknown strain. For *S. mikatae* IFO1815, a potential donor might be a *S. cerevisiae* strain from haplogroup C1b, but its amino acid sequence is similar to IFO1816 (Figure S1). Unfortunately, IFO1815 and IFO1803 strains were not further explored in this study.

### 4.2 Two well differentiated *ORF1* gene sequences define the two *S. cerevisiae* haplogroups

To define the extension of the recombination, we sequenced the *ORF1* region from 36 strains, representative of the most frequent *COX2* haplotypes. Together with previously sequenced *Saccharomyces* strains, we analyzed a total of seventy-two *ORF1* sequences (Table S1). The presence of *ORF1* gene was confirmed for the six available *Saccharomyces* species, indicating its broad dissemination across the genus. *ORF1* length is highly polymorphic among strains and species, mostly due to the differing content of AT-rich tandem repeats and GC clusters (see Supplementary Text, Figure S3), with lengths ranging from 1.3 Kb *(S. cerevisiae* ZA17) to 1.5 Kb *(S. cerevisiae* VRB).

The *ORF1* phylogenetic tree (Figure S3) conflicts with the species tree (Figure 3), indicating the presence of reticulate events driven by a potential HGT from one species to another or the presence of recombinant sequences. To visualize the presence of conflicting data in the alignment, we reconstructed a NN phylogenetic network (Figure 4). Two groups of sequences were visualized in this network: the type I group contains most *Saccharomyces ORF1* sequences, except for sequences from *S. kudriavzevii*. The type II group contained *S. cerevisiae* haplotypes from the *COX2* haplogroup C2 and the sequence of the Far Eastern *S. paradoxus* CECT11152, which was detected as recombinant for the *COX2* gene (Figure S1 and S2). Haplotypes from some *S. cerevisiae,* some European and Far Eastern *S. paradoxus, S. kudriavzevii* strains, as well as some hybrids, were located in an ambiguous position in the phylogenetic network, between the two main *ORF1* groups, suggesting they contained recombinant sequences.

**Figure 4.**
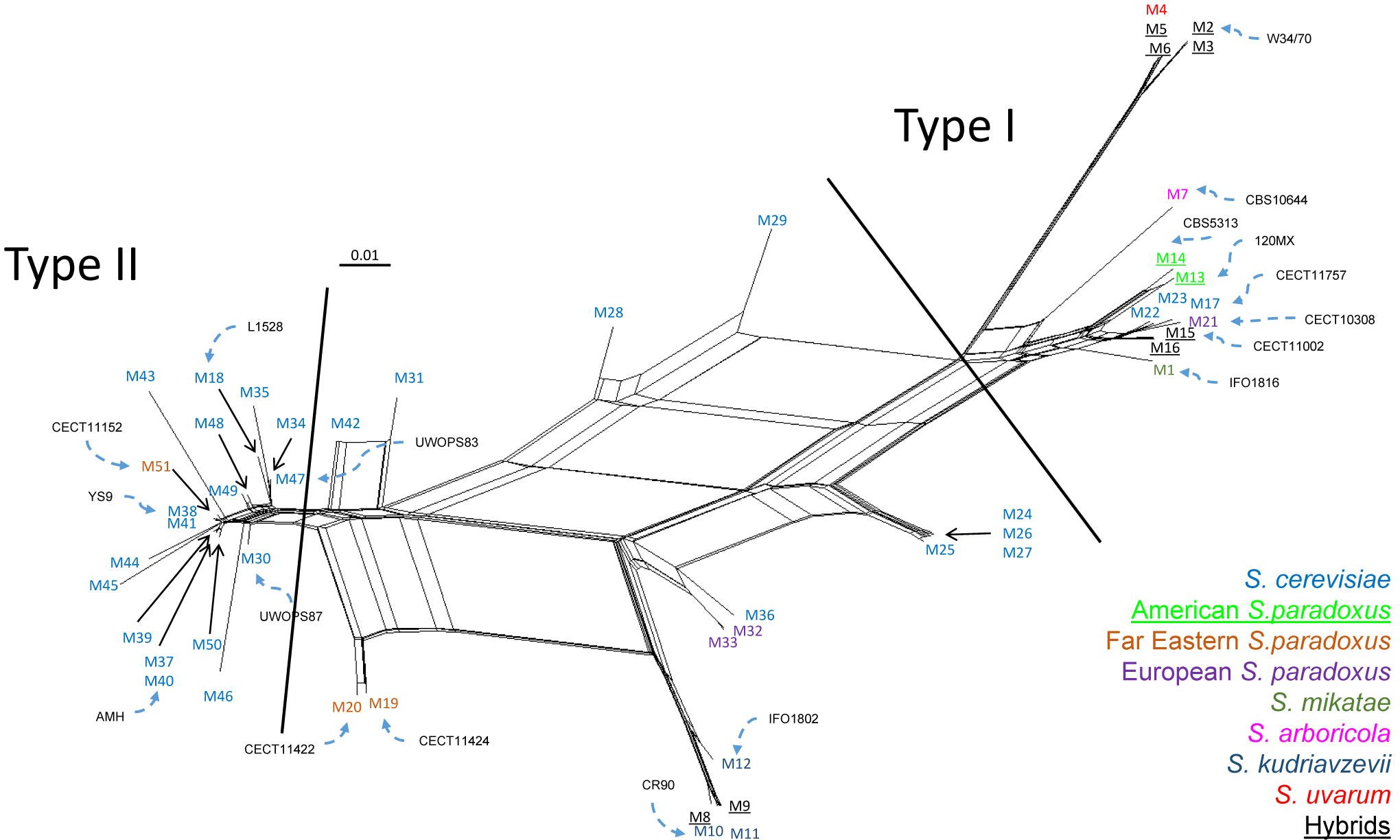
*ORF1* Neighbor-Net phylogenetic network. The phylogenetic network differentiated two groups of *ORF1* sequences, type I and type II. Haplotypes for *ORF1* are represented with M[number] according to Table S1. Haplotype numbers are colored according to the species designation. Scale is given in nucleotide substitution per site. Strain names discussed in the main text are represented and connected to their corresponding *ORF1* haplotype by a dashed blue arrow.

Recombinant analyses, using the *ORF1* sequence of seventy-two representative *Saccharomyces* strains, supported the presence of four partitions (`GARD` reported ΔAICc 143.724 when moved from 3 to 4 partition model, and no improvement, ΔAICc 0, when moved from 4 to 5). Due to the complexity of the data showing different recombinant points, `RDP` and `GARD` disagreed with the location of the second breakpoint (Figure S4 and S5B).

As commented above, *ORF1* sequences from Far Eastern *S. paradoxus* and *S. kudriavzevii* strains were found in an ambiguous position, and closely related for some partitions (Figure S5B,C). Analysis of genetic distance for the different partitions in *S. kudriavzevii* and Far Eastern *S. paradoxus* detected a high identity among *S. kudriavzevii* and Far Eastern *S. paradoxus* (CECT11422 and CECT11424) *ORF1* amino acid sequences for two segments of the alignment (Figure S4 and S5). The genetic distance for the second region was: 1.1% *(S.par* FE-S.kud IFO1802) and 2.4% *(S. par* FE-S.kud EU), much lower compared to genetic distance calculated from nuclear genomes, 13.66% and 13.55%, respectively, and 46.05% *(S. par* FE-S.kud IFO1802) and 47.91% *(S.par* FE-S.kud EU) when compared to the fourth *ORF1* segment (Figure S4 and S5).

Particular attention must be given to the *S. cerevisiae* strains CECT11757 and L1528, representative of the C2xP3 *(S. cerevisiae* C2 × Far East *S. paradoxus*, Figure S1, and S2) recombinant *COX2* group, and which sequence was also contained in hybrids IF6, UvCEG, and S6U. These two strains bear an *ORF1* type I closely related to European CECT10308 *ORF1* (Figure S4 and S5), instead of the *ORF1* sequence observed for Far Eastern *S. paradoxus* CECT11422 or CECT11424. There are two possible explanations: i) a second HGT from European *S. paradoxus* into these strains, which were previously introgressed by Far Eastern *S. paradoxus*, and now showing Far Eastern *S. paradoxus* polymorphisms just at the end of *COX2;* ii) or a patchy-tachy model (Sun *et al*. 2011) in the *COX2* 3’end region driving homoplasic mutations, where the *S. cerevisiae* strains in C2xP3 group, now contain similar polymorphisms than Far Eastern *S. paradoxus* strains. In the case of L1528, the recombination track was prematurely resolved, containing a chimeric *ORF1* type I × type II (Figure S4 and S5).

### 4.3 *COX3* supports the HGT/introgression in the *COX2-ORF1* region and noncommon introgressions for the American *S. paradoxus*

To improve the species assignment based on a mitochondrial gene, we analyzed the *COX3* sequence of the selected seventy-two *Saccharomyces* strains (Table S1). The lack of introns and overlapping homing endonucleases make *COX3* a good candidate for the species assignment.

The *COX3* NJ phylogenetic tree was mostly congruent with the species tree (Figure 3 and S6), except for the American *S. paradoxus*, which strains were enclosed with *S. cerevisiae* strains. In addition, the position of *S. mikatae* was not well resolved. The *COX3* MJ phylogenetic network assigned species by haplogroups (Figure 5). We located *S. eubayanus* haplogroup with W34/70 haplotype, as demonstrated by Baker *et al*. (2015) and Okuno *et al*. (2016). We found a high number of nucleotide substitutions in the *S. cerevisiae COX3* sequences retrieved from the *Saccharomyces* Genome Resequencing Project (SGRP) (H4, H9-11), indicative of assembling errors for that particular gene; nevertheless, the topologies of the MJ network and the NJ tree were not affected.

**Figure 5.**
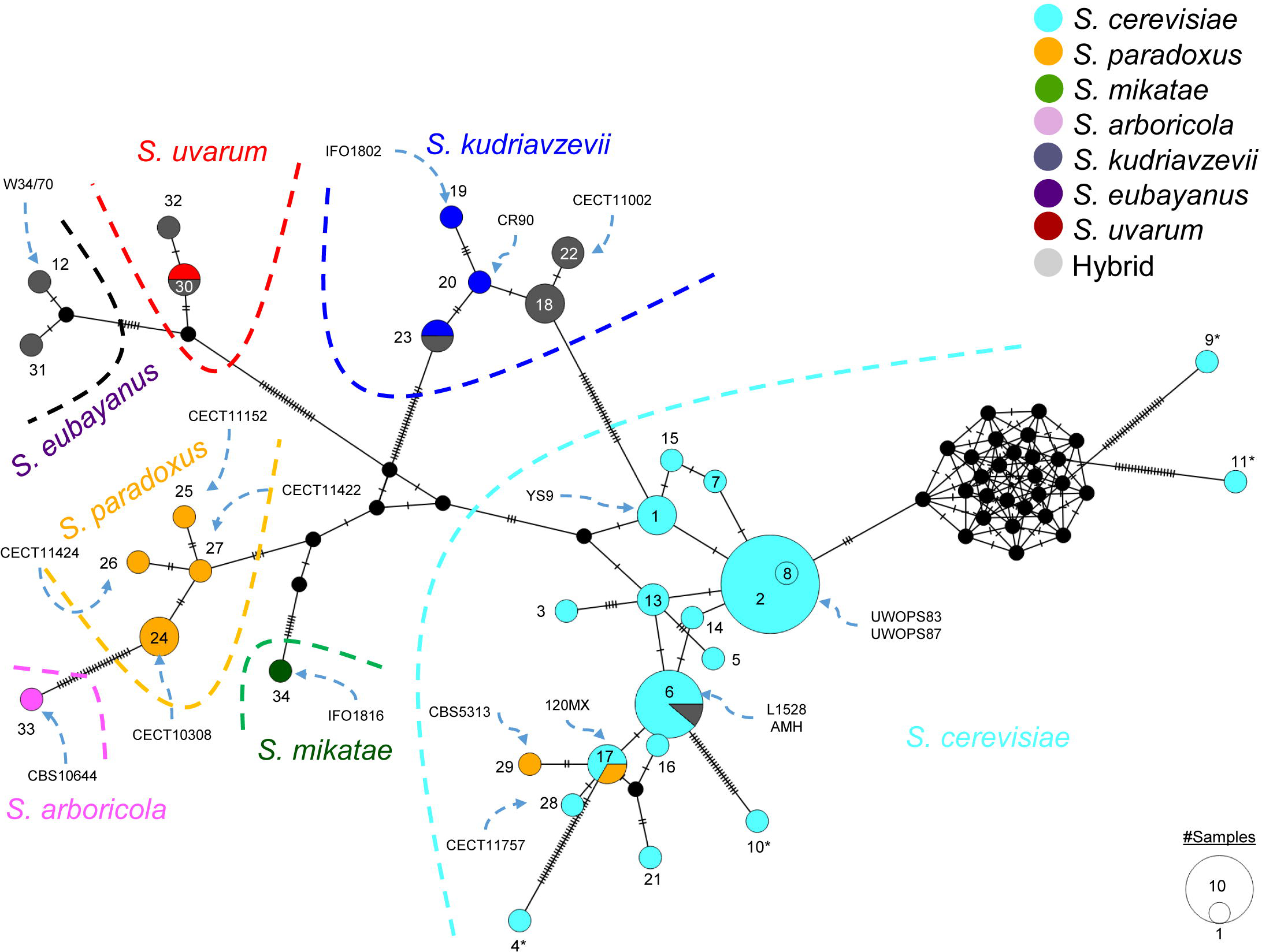
*COX3* Median-Joining phylogenetic network. Thirty-four *COX3* haplotypes (Table S1) were represented by circles. Circle size is scaled according to the number of sequences in a haplotype. Pie charts show the frequency of each species in that particular haplotype. Asterisks indicate haplotype sequences with nucleotide errors. Number of mutations from one haplotype to another are indicated by lines in the edges connecting haplotypes. Strain names discussed in the main text are represented and connected to their corresponding *COX3* haplotype by a dashed blue arrow.

The *S. paradoxus* CECT11152 (syn. IFO1804), which was enclosed with *S. cerevisiae* strains for the *COX2* and the *ORF1* gene (Figure 2 and 4), was within *S. paradoxus COX3* haplogroup (Figure 5 and S6). This result confirms that the HGT event was localized between the 3’ end *COX2* and the *ORF1* region (Figure S2B). A similar conclusion is reached for *S. cerevisiae* × *S. kudriavzevii* hybrids (CECT11002, CECT11011, and CECT1990), where the flanking regions of *ORF1* are from a *S. kudriavzevii* donor. The two representative *S. cerevisiae* strains (L1528 and CECT11757), from the C2xP3 recombinant *COX2 (S. cerevisiae* × Far Eastern *S. paradoxus*, Figure 2), and containing all or most of the *ORF1* sequence from a *S. paradoxus* donor, were enclosed with *S. cerevisiae COX3* sequences, supporting the detected HGT for the *COX2-ORF1* region (Figure 5 and S6).

Our two representatives from the American *S. paradoxus*, 120MX and CBS5313, displaying an identical 5’ end *COX2* segment sequence to *S. cerevisiae* sequences (Figure S1A), also had a *COX3* sequence closely related to the *S. cerevisiae* haplogroup. The *COX3* haplotype 17 of 120MX was identical to *S. cerevisiae COX3* sequences (Figure 5 and S6). This result suggests that the American *S. paradoxus* likely inherited the flanking regions of *ORF1* from *S. cerevisiae*. Inspection of complete American *S. paradoxus* mitochondrial genome will shed light about the extension of the introgressed region.

### 4.4 Recombinant sites are close to the starting *ORF1* CDS, regions of integration of GC clusters and A+T-rich sequences

We reanalyzed the recombinant sites using a concatenated alignment of the three genes studied here (~2.3 Kb). This analysis detected additional recombinant points, that together with the previous sites using individual genes made a total of 14 recombinant sites (Figure 6). In addition to recombinant points found at the beginning of the *ORF1* CDS, some recombination breakpoints were located close to A+T-rich sequences or regions were GC clusters can be integrated (see Supplementary text, Figure 6). Two of the recombination points were just on the beginning of both *ORF1* LAGLIDADG domains (Figure 6). At least one of the recombinant events involved some of the haplotypes located in the *ORF1* phylogeny in-between the two main types (Figure 4, 6, and S5). Recombinations were detected between Type I × Type I, Type I × Type II and Type II × Type II (Figure S6 and S7). It was slightly significant the finding of eleven of the fourteen recombinant sites in the surroundings of regions with the presence of recombinogenic sequences, such as the beginning of the putative homing endonuclease *ORF1*, regions of integration of GC clusters and A+T tandem repeats (X^2^ test p-values < 0.03251), suggesting these genetic elements are responsible of the initiation of most of the recombination events.

**Figure 6.**
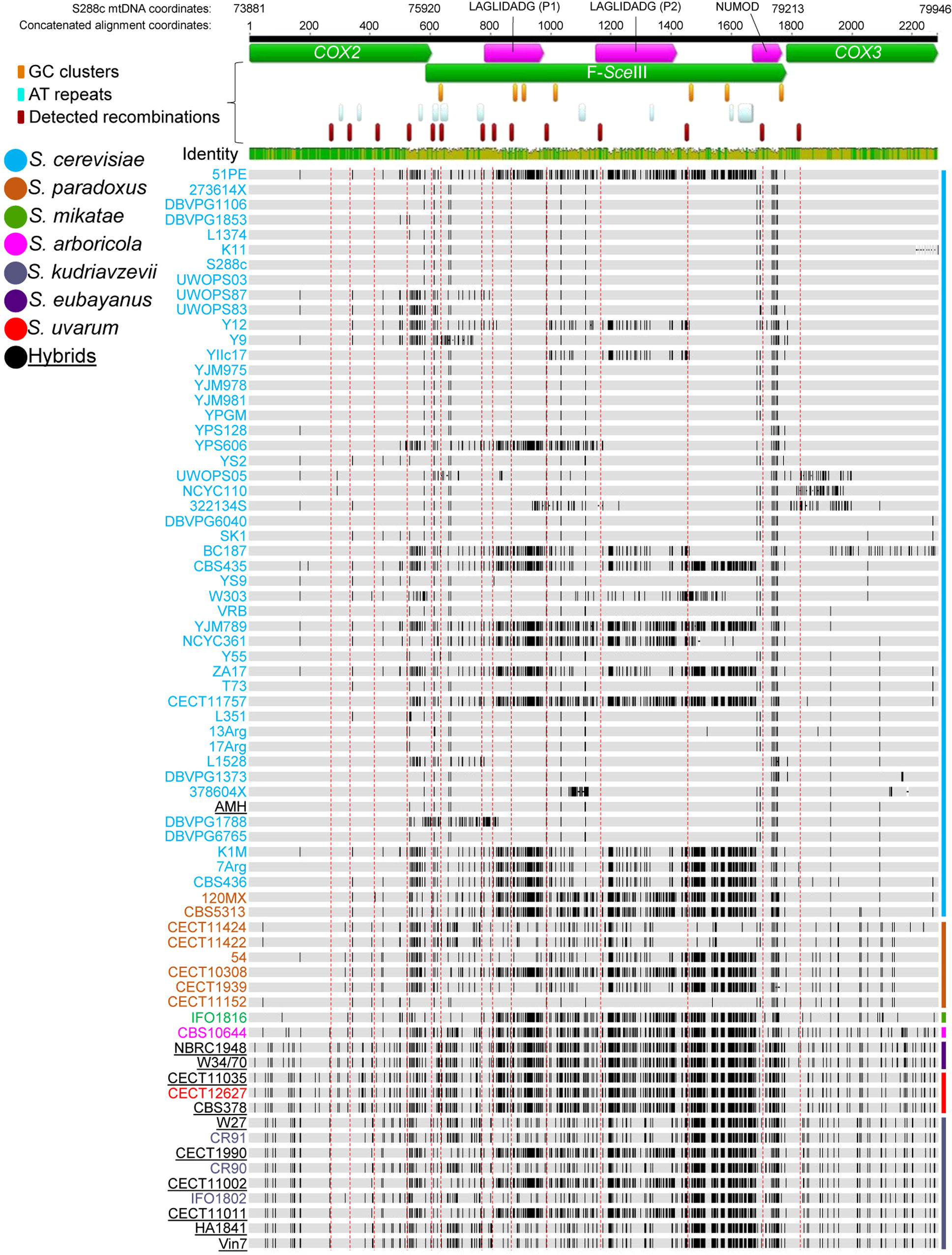
*COX2-ORF1-COX3* concatenated alignment. A representation of the concatenated alignment of *COX2-ORF1-COX3* region (~2.3 Kb) was shown. Genes were represented by green. The *ORF1* LAGLIDADG and NUMOD domains were represented in pink. Aminoacid conservation of the domains are shown by WebLogo representations in Figure S9. A completed representation of the aminoacids of the overlapped genes, *COX2* and *ORF1*, together with the annotation of alpha helixes and beta sheets of the domains can be found in Figure S9 and S10. Detected A+T tandem repeats (Table S4), GC Clusters (Figure S11, S12), and recombination points are represented according to the legend. Identity graph represents the percentage of nucleotide conservation for each position. Polymorphic sites are represented by black lines. Dashed red lines are the detected recombination points for helping during visualization. Sequences were clustered based on the *COX3* gene sequence. Color lines in the right of the sequences represents the species *COX3* sequence (Figure 5) according to the legend. Strain names were colored according to the legend.

A visualization of the polymorphisms of the concatenate alignment clearly demonstrated that recombinant track is resolved differentially depending of the strain. But in all cases the recombinant track is resolved before getting the *COX3* region (Figure 6), which is ~3.4 Kb apart from the last *ORF1* nucleotide explored here. It is important to note that *COX3* in *S. paradoxus* strains is in a different location in the mitochondrial genome, approximately ~30 Kb apart (Procházka *et al*. 2012).

### 4.5 All *S. cerevisiae COX2* haplogroups are worldwide distributed

The recombination located in *COX2* gene makes this gene a good candidate to differentiate closely related strains. Those strains sharing a similar recombination are expected to share a similar ancestor. To describe the phylogeography of *S. cerevisiae,* we explored the polymorphic *COX2* gene of 418 *S. cerevisiae* strains from 7 continents, isolated from both human-associated (baking, beer, clinical, laboratory, sake, wine and traditional alcoholic beverages) and wild environments (Figure 1, Table S1).

An association study among *S. cerevisiae* strain origins and their haplogroup distribution was performed (Figure S7A). Clinical and wine samples were significantly associated with haplogroup C2 (χ^2^ test p-values 9.9 × 10^−5^ and 3 × 10^−4^, respectively). We applied a similar approach to infer the distribution of haplogroups taking into account their geographic origins (Figure S7B), revealing that European *S. cerevisiae* strains were highly associated with Haplogroup C2 (χ^2^ test p-value 9.9 × 10^−5^). No significant distribution bias was observed in the remaining strains, possibly due to the low number of wild isolates in our study. For example, beer strains were included in both haplogroups with no clear association, maybe due to the presence of multiple beer lineages (Gonçalves *et al*. 2016; Gallone *et al*. 2016).

## 5. Discussion

### 5.1 Wine *S. cerevisiae* domestication bottleneck fixed closely related *COX2* haplotypes

Two independent *S. cerevisiae* domestication events from wild isolates have been inferred (Legras *et al*. 2007), one for sake and another for wine strains (Fay and Benavides 2005; Liti *et al*. 2009; Schacherer *et al*. 2009; Almeida *et al*. 2015). Indeed, *S. cerevisiae* wine domestication is attributed to be originated in the Near East (Fay and Benavides 2005; Liti *et al*. 2009) probably from the wild *S. cerevisiae* stock from Mediterranean oak (Almeida *et al*. 2015). A high frequency of European wine strains within haplogroup C2 was detected, suggesting that the bottleneck during the domestication to winemaking fixed *COX2* variants from haplogroup C2. This data might indicate an impact of the haplogroup C2 on the winemaking process. If we are correct in our hypothesis, we would expect strains with C2 haplotype to perform better under winemaking conditions. Previous works have demonstrated the fitness effect driven by transferring the mitochondrial genome of one *S. cerevisiae* strain to another *S. cerevisiae* strain with a different haplotype (Zeyl *et al*. 2005; Paliwal *et al*. 2014), supporting the effect of different mitochondrial genomes in the fermentation performance. It might be interesting to see whether the complete mitochondrial genome clusters most of wine/European strains, and it has a positive impact during wine fermentations compared to mitochondrial genomes from non-wine isolates.

Despite most of the wine isolates fixed the *COX2* C2 haplotype, the observation of wine strains within *COX2* haplogroup C1a and C1b, such as EC1118, and non-wine strains enclosed with C2 haplogroup, might suggest recent gene flow among the different lineages in wild environments. This scenario has been suggested by population genomic analysis of wild and domesticated lineages (Almeida *et al*. 2015). It is also supported by the observation of reintroduced wine isolates into wild environments, such as oak trees, close to vineyards in non-European regions (Hyma and Fay 2013), and the presence of admixture strains between wine and non-wine lineages (Liti *et al*. 2009; Schacherer *et al*. 2009; Gallone *et al*. 2016). Most clinical samples are derived from wine isolates, as the mitochondrial haplogroup indicate. This is in agreement with previous results of the 100 *S. cerevisiae* project (Strope *et al*. 2015; Wolters *et al*. 2015).

### 5.2 Most industrial hybrids contain an introgressed or a non-cerevisiae mitochondrial DNA

Hybrids can fix the mitochondrial genome of one of the two parents, or a recombinant version, reaching the homoplasmic state in few generations (Strathern *et al*. 1981, Berger and Yaffe 2000). Interestingly, most of our hybrids, except AMH, have inherited a non-*cerevisiae* (Peris *et al*. 2012a; Peris *et al*. 2012b; Peris *et al*. 2014; Pérez-Través *et al*. 2014) or an introgressed mitochondrial genome (this study, Peris *et al*. 2014), which might suggest a better fermentative performance of this haplotype to industrial conditions where this hybrids were selected, which usually are characterized by low fermentation temperatures (Belloch *et al*. 2008, Peris *et al*. 2012a, Gibson *et al*. 2013). It is also clear how in the case of the dietary supplement and wine hybrids *S. cerevisiae* × *S. kudriavzevii* (IF6 and UvCEG), and a winemaking hybrid *S. cerevisiae* × *S. uvarum* (S6U) inherited a mitochondria genome from a *S. cerevisiae* strain donor, closely related to wine strains isolated from Europe, South America and Africa, containing an introgressed mitochondria. Before the hybridization event between *S. cerevisiae* with other *Saccharomyces* species, *S. cerevisiae* mitochondria was introgressed with a European *S. paradoxus* (Figure 7). Introgressions occurring before hybridization might be also a potential scenario for some other hybrids, such as *S. cerevisiae* × *S. kudriavzevii* hybrids which showed introgressions from European *S. paradoxus* (Peris *et al*. 2012a) (Figure 7) or introgressions from European *S. uvarum* into mitochondria of Holarctic strains of *S. eubayanus* (Peris *et al*. 2016), the close relatives of the parental donor of *S. cerevisiae* × *S. eubayanus* hybrids.

**Figure 7.**
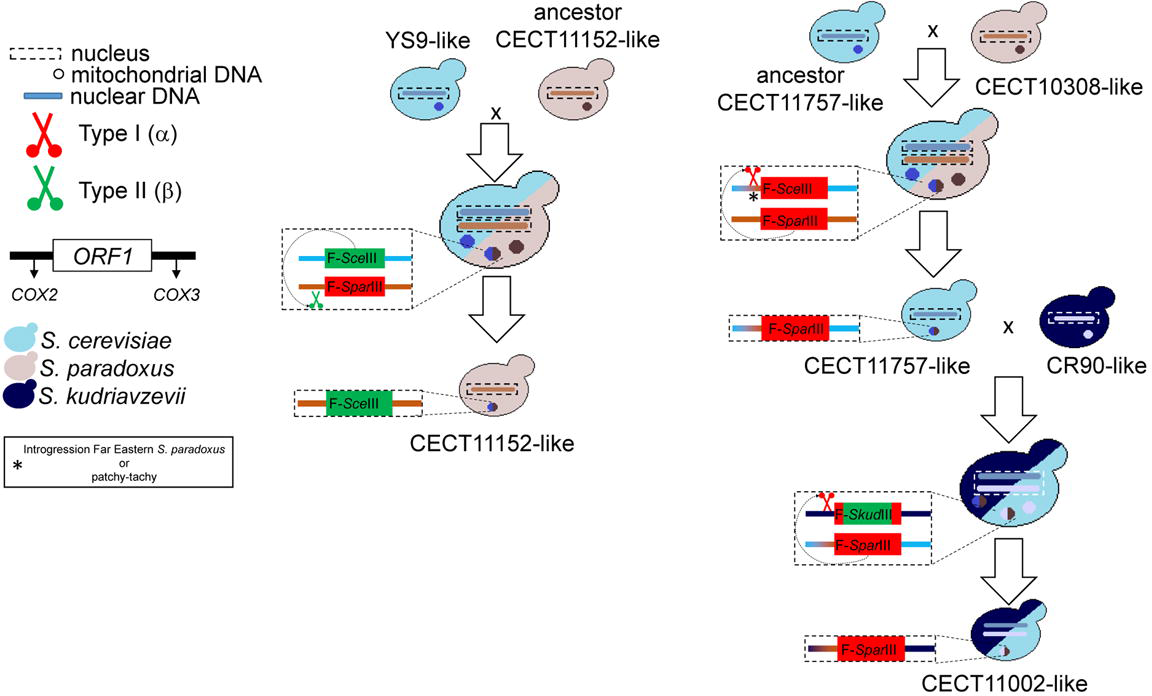
Summary *ORF1* invasion in a wild isolate and a hybrid. A summary of two introgression events potentially driven by the invasion of the homing endonuclease *ORF1* was represented for the Far East *S. paradoxus* CECT11152, and the *S. cerevisiae* × *S. kudriavzevii* hybrid CECT11002. The two *ORF1* main types, type I and type II, are represented by red and green, respectively. The *COX2* and *COX3* are represented to highlight the mitochondrial background. When nuclear genomes in the hybrid state are double, it represents that nucleus do not fuse. In the case of the *S. cerevisiae* × *S. kudriavzevii* hybrid just one nuclear genome is represented because it is well know the presence of chromosomes from both parents (Peris *et al*. 2012c). The scissor symbol represents the homing endonuclease protein. Small circles represent the mitochondrial DNA. When double color is shown in the mitochondrial symbol represent a mixture of both mitochondrial DNA that might originate the recombinant versions as demonstrated in this study. For more hypothesized complex models visit the Figure S8.

Previous work has demonstrated the impact in respiration levels of the inheritance of different mitochondrial genomes, *S. cerevisiae* or *S. uvarum*, by hybrids (Albertin *et al*. 2013), highlighting the importance of taking into account the fixation of the mtDNA during the generation of artificial hybrids for specific industrial processes. However, we still do not understand the impact of the introgression described in this study in the fitness of these hybrids. In addition, the impact of the inheritance of other mitochondrial genomes, such as *S. kudriavzevii*, in the adaptability of hybrids is still unknown.

### 5.3 Mitochondrial introgressions as evidence of ancestral hybridization events in wild environments

*Saccharomyces* double and triple hybrids have been isolated from many different industrial conditions, such as “ale” and lager beer, wine, cider, dietary supplements (Casaregola *et al*. 2001; Barros Lopes *et al*. 2002; González *et al*. 2008; Peris *et al*. 2012a; Pérez-Través *et al*. 2014) and clinical samples (Peris *et al*. 2012a). Hybrids were thought not to occur in the wild environment, suggesting that hybridization is only limited in artificial conditions where hybrids are better adapted to those stressful environments (Belloch *et al*. 2008) by the acquisition of beneficial traits from the parents (Peris *et al*. 2012c; Gibson *et al*. 2013). To accommodate the idea of non-hybridization in wild environment, HGT between *S. cerevisiae* and *S. paradoxus* has been suggested to be mediated by the formation of heterokaryons (Wu *et al*. 2015). These heterokaryons are the result of pseudohyphae formation by *Saccharomyces* strains allowing the contact for those strains found in sympatric association (Wu *et al*. 2015). However, a complete hybridization with a rapid loss of one of the parent genomes by outcrossings with the non-hybridized sibling strains might be another potential scenario. The diversity of *S. cerevisiae* × *S. kudriavzevii* hybrids (Peris *et al*. 2012a), most of them with lower *S. kudriavzevii* genome content, suggests a rapid loss of *S. kudriavzevii* genes after hybridization while keeping traits important for low temperature fermentations (Peris *et al*. 2012c). In addition, several introgressions involving most *Saccharomyces* species suggest some gene flow (Liti *et al*. 2005; Muller and McCusker 2009; Almeida *et al*. 2014). The hypothesis of hybridization in wild environment has been recently highly supported by the isolation of hybrids *S. cerevisiae* × *S. paradoxus* from Brazil (Barbosa *et al*. 2016).

### 5.4 Introgressions might influence diversification

The two clearest mitochondrial introgressions were modeled in Figure 7. All potential transfers detected in our study are summarized in Figure S8 which is a starting point to model mitochondrial introgressions in *Saccharomyces* genus. Some particular *Saccharomyces* lineages have some degree of introgression, such as Asia B *S. kudriavzevii*, Asia A and European *S. kudriavzevii*, and *S. mikatae* IFO1815.

The most interesting introgression was detected in all American *S. paradoxus* from populations B and C. These strains have a *COX2* sequence with a closely related 5’ end to *S. cerevisiae* from haplogroup C1, and the two selected representative *S. paradoxus* strains (120MX and CBS5313) shared a *COX3* sequence with *S. cerevisiae* strains. Recent evidence of phylogenetic tree incongruence among *S. cerevisiae* and American population B *S. paradoxus* YPS138 (Leducq *et al*. 2014) strain was also detected (Wu *et al*. 2015), supporting our hypothesis. The increase of new isolates resulting from the improvement of isolation methods (Sampaio and Gonçalves 2008; Sylvester *et al*. 2015), along with the application of new methods for assembling mitochondrial genome will shed light about other American *S. paradoxus* mitochondrial inheritance and the extension of the mitochondrial introgression detected here.

The presence of mitochondrial introgressions raises the question: has mitochondrial introgression influenced the phenotypes and the diversification of *Saccharomyces* species? The empirical data commented in previous sections might support mitochondrial has an influence in the diversification of yeast species. In this way, the accommodation of the nuclear genome to this new mitochondrial genome might drive the diversification of American *S. paradoxus* from the other *S. paradoxus* lineages. An interesting question that will require experimental demonstration and more data from additional strains.

### 5.5 Recombinations in the *COX2-ORF1* region might be mediated by *ORF1*, regions of integration of GC clusters and/or A+T tandem repeats

Evidence of intraspecific mitochondrial recombination among *S. cerevisiae* strains has been shown (Dujon *et al*. 1974; Nunnari *et al*. 1997; Berger and Yaffe 2000). Indeed, a complete DNA recombination map in *S. cerevisiae* has been recently drawn (Fritsch *et al*. 2014). Mitochondrial recombination is initiated by a double-strand break (DSB) generated and resolved by mainly four nuclear encoded proteins (Lockshon *et al*. 1995; Ling and Shibata 2002; Ling *et al*. 2007; Ling *et al*. 2013) and it can be facilitated by the mitochondrial genome architecture, presence of GC clusters, or by the mobility of mitochondrial elements, such as introns or homing endonucleases (Dieckmann and Gandy 1987; Séraphin *et al*. 1987; Yang *et al*. 1998). We found that two *COX2 S. cerevisiae* and *S. paradoxus* groups of sequences are mainly driven by the presence of two highly divergent *ORF1* gene sequences, type I and type II. The presence of two clear *COX2* groups might be the result of the *ORF1* invasion. The origin of this *ORF1* is still unclear but the presence of *ORF1* type I in most of *Saccharomyces* species might suggest this is the ancestral *Saccharomyces ORF1* homing endonuclease, and the type II might be from *non-Saccharomyces* origin.

The clearest example of *ORF1* transfer was between Far Eastern *S. paradoxus* CECT11152 (syn. IFO1804) and *S. cerevisiae* (Figure 7), which is supported by an additional detected HGT of the GC cluster GC42 between *S. cerevisiae* and the Far Eastern *S. paradoxus* CECT11152 (Wu and Hao 2015), suggesting that both transfers likely occurred at the same time. For this reason, we propose that *ORF1* is likely a functional homing endonuclease similar to other *S. cerevisiae* free-standing homing endonucleases, such as *ORF3 (ENS2)* (Séraphin *et al*. 1987; Nakagawa *et al*. 1992). Following nomenclature rules (Belfort and Roberts 1997), we suggest renaming *ORF1* to F-*Sce*IIIα or F-*Sce*IIIβ for *S. cerevisiae*, and replacing the last two letters of *Sce* with the corresponding first two letters for each species *ORF1*, where “F” stands for freestanding homing endonuclease, “S” for *Saccharomyces*, “ce” for *cerevisiae*, and III because F-*Sce*I and F-*Sce*II are designated for Endo.*Sce*I *(ENS2, ORF3)* and HO endo, respectively. The type I and type II group membership is designated by using a suffix α and β, respectively.

A recent study, where fourteen *S. cerevisiae* and *S. paradoxus* mitochondrial genomes were sequenced, found HGTs between those two species mainly facilitated by GC clusters (Wu *et al*. 2015; Wu and Hao 2015). Our results showed recombination breakpoints close to regions of integration of GC clusters and A+T tandem repeats in wild *Saccharomyces* strains, supporting that these genomic elements are facilitating mitochondrial recombination events. All these three elements, regions of integration of GC clusters, A+T tandem repeats and homing endonucleases might be responsible of facilitating recombination in mitochondria.

## 6. Conclusions

Reticulate evolutionary events, such as recombination, introgression and horizontal gene transfers, are more common than previously thought, and might have an ecological impact in the species undergoing such events. Although, the mechanisms in yeasts are still unclear and the effects to the phenotype must be clarified, it appears that homing endonucleases, such as F-*Sce*III, and other genetic elements play an important role in the transmission of genetic material among different species, with important outcomes in the generation of yeast diversity.

## 7. Acknowledgments

We thank to C.P. Kurtzman to provide us the *S. mikatae* NRRL Y-27342. We thank Chris Todd Hittinger and William G. Alexander for critical comments on the manuscript. This work was supported by Spanish Government grant (AGL2009-12673-C02-02) and Generalitat Valenciana grants (PROMETEUS and ACOMP/2012) to EB, and from the Spanish Government FEDER (AGL2012-39937-C02-01) and Generalitat Valenciana (PROMETEOII/2014/042) to AQ. DP acknowledges to the Spanish Government for its Ministerio de Ciencia e Innovacion (MICINN) FPI fellowship. AA received PROMEP Fellowship from SEP, Mexican government. LP acknowledges to CSIC and the Spanish Ministry of Education and Science (MEC) for an I3P fellowship. SO acknowledges to MEC for the postdoctoral research contract.

## 9. Figure Legends

**Table S1. Strain information.**

**Table S2. Primer pairs used in this study.**

**Table S3. Summary statistics for all Saccharomyces and each species using *COX2*.**

**Table S4. Distribution of AT tandem repeats**.

**Figure S1. *COX2* polymorphic sites**. Variable *COX2* nucleotide positions, and discriminant aminoacids (in the right hand of the figure) encoded by highlighted red nucleotides, are shown. Haplotype numbers are colored according to the species designation, according to Figure 2 and the legend. Numbers above polymorphic sites represent their position in the alignment. Black bars in the bottom highlights the *COX2* regions used for reconstructing the phylogenetic networks in Figure S2. Haplogroup (HapG) designation and the number of sequences (#seq) for each haplotype are shown in the left hand. The position in the alignment of the first nucleotide for each codon, encoding the represented discriminant aminoacids, are shown above the aminoacid sequence. Sequences were colored according to their similarity as inferred from Figure S2B. Strain names discussed in the main text are represented in the right hand of their corresponding haplotype.

**Figure S2. *COX2* Neighbor-Net phylogenetic network for each *COX2* segment**. Reconstructed neighbor-net phylogenetic networks using the *COX2* 5’ end, and 3’ end segments are represented in A) and B), respectively. Haplotypes were colored according to each species designation. Haplotypes exclusively from hybrids are colored in grey. Scale is given in nucleotide substitution per site. Strain names discussed in the main text are represented and connected to their corresponding haplotype by a dashed blue arrow.

**Figure S3. *ORF1* Neighbor-Joining phylogenetic tree**. A NJ phylogenetic tree was reconstructed for the *ORF1* sequence alignment. Strains were colored according to the species designation. Bootstrap values above 50 are given for each branch. A red star represents a potential active *ORF1* based on the absence of GC clusters, which could disrupt the coding sequence, and premature stop codons. Type I and Type II *ORF1* sequences, detected as non-recombinant (represented by dots), were colored in red and green, respectively. GC clusters in the *ORF1* sequence were represented by squares and the number represents the position in the *ORF1* alignment (see Figure 6). An arrow indicates that a particular GC cluster was inverted to infer the GC cluster family. Square colors represent GC cluster similarity according to Figure S11, and the NJ trees from Figure S12.

**Figure S4. *ORF1* aminoacid polymorphic sites**. Variable *ORF1* aminoacid positions among haplotypes. Haplotype number are colored according to the species designation, as in Figure 2. Alignment aminoacid positions for each polymorphic site are also shown. The *COX2* Haplogroup designation is shown. Sequences were colored according to their similarity as inferred from Figure S5. Symbols * and ^ represent type I and type II *ORF1* sequences, respectively. Regions colored in yellow are sequences from unknown source. Lines indicate the sites corresponding to each alignment partition to reconstruct the *ORF1* phylogenetic networks by segments (Figure S5).

**Figure S5. *ORF1* NN phylogenetic networks by segments**. NN phylogenetic networks. for each *ORF1* alignment partitions, inferred by `GARD` and `RDP`, are shown in A), B), C) and D). A) shows the NN phylogenetic network corresponding to the region from *COX2* 3’ end (Figure S1) to the position 246 of the *ORF1* alignment. Nucleotide positions from *ORF1* alignment are shown in figures B-D. Scale bar are given in nucleotide substitution per site. Type I and Type II *ORF1* sequences, detected as non-recombinant, are represented by red and green circles, respectively. Sequences are colored to each species designation. In figure B) some S. *cerevisiae* sequences are grouped in a I -> II group, indicating that the region used for inferring that NN phylonetwork contains a recombination point for those particular strains driving to the ambiguous position. Asterisks highlight sequences which clustering changed from one *ORF1* type to another due to its recombinant character.

**Figure S6. *COX3* NJ phylogenetic tree**. A *COX3* Neighbor-Joining phylogenetic tree is shown. Strains are colored according to the species designation. Bootstrap values above 50 are given for each branch. Scale is given in nucleotide substitution per site.

**Figure S7. *S. cerevisiae COX2* MJ networks**. *S. cerevisiae COX2* MJ networks were represented in A) and B) where haplotype pie charts were colored according to the isolation source and continent of isolation, respectively. Circle sizes represent the number of sequences in a haplotype and is scaled according to the legend. Number of mutations from one haplotype to another are indicated by lines in the edges connecting the haplotypes. A table showing the strain percentage distribution in each haplogroup or recombinant group by isolation source or continent is also displayed in A) and B), respectively.

**Figure S8. A summary model of potential *COX2-ORF1* introgression in wild environments and during domestication**. A starting model of potential introgressions in *Saccharomyces* is described. Black, light blue, orange, green, dark blue, pink, red and purple yeasts represent the yeast ancestors, *S. cerevisiae, S. paradoxus, S. mikatae, S. kudriavzevii, S. arboricola, S. uvarum* and *S. eubayanus* yeasts, respectively. Small circles inside yeasts represent the mtDNA, which is colored according to the species designation. A *S. paradoxus* yeast with a blue small circle indicates a potential inheritance of mtDNA from *S. cerevisiae*. Red, green or red/green boxes represent type I, type II or recombinant *ORF1* sequences, respectively. A question tag indicates dubious scenario, unknown *ORF1* or unknown mtDNA sequence due to the absence of *COX3* sequence to support the mtDNA inheritance.

**Figure S9. Weblogo representation of LAGLIDADG and NUMOD domains**. Weblogo (<u>http://weblogo.berkeley.edu/</u>) of the three homing endonuclease domains for *Saccharomyces ORF1* genes plus *SasefMp08* from *Kazchastania servazii*, and *ORF1* and *ORF3* from *Cyberlindnera (Williopsis) saturnus* var. *suaveolens* are represented. Cylinders and arrows represent α-helix and β-sheet, as previously described (Dalgaard *et al*. 1997). Asterisk symbol indicates the codons for those aminoacids under purifying selection detected by DataMonkey.

**Figure S10. *COX2* and *ORF1* aminoacid alignment (Supplementary File Figure S10)**. The complete *COX2* and *ORF1* aminoacid alignments are shown. Protein secondary structure and domains are indicated for *ORF1*. Jalview also display the alignment quality (based on BLOSUM 62), physicochemical conservation calculated according to Livingstone and Barton (Livingstone and Barton GJ 1993), and consensus sequence.

**Figure S11. GC clusters from M1 family insertion regions**. The flanking regions where the seven GC clusters were found are shown. In the case of GC cluster 6 and 7 a consensus sequence is shown and bars show the percentage of sequences showing a particular nucleotide substitution. An arrow indicates that the GC cluster sequence was inverted to annotate its structure. Question marks indicate an unknown GC cluster structure.

**Figure S12. GC cluster 6 and 7 Neighbor-Joining phylogenetic trees**. To classify the GC cluster according to sequence similarity we reconstructed the NJ phylogenetic tree of GC cluster 6 and 7. This classification is shown by numbers in Figure S3. Scale bars represent number of substitutions per site.

## References

Adler D, Murdoch D. 2009. rgl: 3D visualization device system (Open GL).

Albertin W, da Silva T, Rigoulet M, Salin B, Masneuf-Pomarede I, et al. 2013. The mitochondrial genome impacts respiration but not fermentation in interspecific *Saccharomyces* hybrids. PLoS ONE. 8:e75121

Almeida P, Gonçalves C, Teixeira S, Libkind D, Bontrager M et al. 2014. A Gondwanan imprint on global diversity and domestication of wine and cider yeast *Saccharomyces uvarum*. Nat Commun. 5.

Almeida P, Barbosa R, Zalar P et al. 2015. A population genomics insight into the Mediterranean origins of wine yeast domestication. Mol Ecol. 24:5412–5427.

Altschul SF, Madden TL, Schaffer AA, Zhang J, Zhang Z, Miller W, Lipman DJ. 1997. Gapped BLAST and PSI-BLAST: a new generation of protein database search programs. Nucl Acids Res. 25:3389–3402.

Andersson JO. 2009. Gene transfer and diversification of microbial Eukaryotes. Annu Rev Microbiol. 63:177–193.

Badotti F, Vilaça ST, Arias A, Rosa CA, Barrio E. 2013. Two interbreeding populations of *Saccharomyces cerevisiae* strains coexist in cachaça fermentations from Brazil. FEMS Yeast Res. 14:289–301.

Baker E, Wang B, Bellora N, Peris D, Hulfachor AB, Koshalek JA, Adams M, Libkind D, Hittinger CT. 2015. The genome sequence of *Saccharomyces eubayanus* and the domestication of lager-brewing yeasts. Mol Biol Evol. 32:2818–2831.

Bapteste E, van Iersel L, Janke A et al. 2013. Networks: expanding evolutionary thinking. 29:439–441.

Barbosa R, Almeida P, Safar SVB et al. 2016. Evidence of natural hybridization in Brazilian wild lineages of *Saccharomyces cerevisiae*. Genome Biol Evol. 8:317–329.

Barros Lopes M, Bellon JR, Shirley NJ, Ganter PF. 2002. Evidence for multiple interspecific hybridization in *Saccharomyces sensu stricto* species. FEMS Yeast Res. 1:323–331.

Basse CW. 2010. Mitochondrial inheritance in fungi. Curr Opin Microbiol. 13:712–719.

Belfort M, Roberts RJ. 1997. Homing endonucleases: keeping the house in order. Nucl Acids Res. 25:3379–3388.

Belloch C, Querol A, Garcia MD, Barrio E. 2000. Phylogeny of the genus *Kluyveromyces* inferred from the mitochondrial cytochrome-c oxidase II gene. Int J Syst Evol Microbiol. 50:405–416.

Belloch C, Orlic S, Barrio E, Querol A. 2008. Fermentative stress adaptation of hybrids within the *Saccharomyces sensu stricto* complex. Int J Food Microbiol. 122:188–195.

Berger KH, Yaffe MP. 2000. Mitochondrial DNA inheritance in *Saccharomyces cerevisiae*. Trends Microbiol. 8:508–513.

Birky CW, Arlene RA, Rosemary D, Maureen C. 1982. Mitochondrial transmission genetics: replication, recombination, and segregation of mitochondrial DNA and its inheritance in crosses. Mitochondrial Genes. Cold Spring Harbor Monograph Archive.

Bordonné R, Dirheimer G, Martin RP. 1988. Expression of the *oxi1* and maturase-related *RF1* genes in yeast mitochondria. Curr Genet. 13:227–233.

Borneman AR, Pretorius IS. 2015. Genomic insights into the *Saccharomyces sensu stricto* complex. Genetics. 199:281–291.

Boynton PJ, Greig D. 2014. The ecology and evolution of non-domesticated *Saccharomyces* species. Yeast. 31:449–462.

Burt A, Koufopanou V. 2004. Homing endonuclease genes: the rise and fall and rise again of a selfish element. Curr Opin Genet Dev. 14:609–615.

Casaregola S, Nguyen HV, Lapathitis G, Kotyk A, Gaillardin C. 2001. Analysis of the constitution of the beer yeast genome by PCR, sequencing and subtelomeric sequence hybridization. Int J Syst Evol Microbiol. 51:1607–1618.

Chou JY, Hung YS, Lin KH, Lee HY, Leu JY. 2010. Multiple molecular mechanisms cause reproductive isolation between three yeast species. PLoS Biol. 8:e1000432

Colleaux L, d’Auriol L, Betermier M, Cottarel G, Jacquier A, Galibert F, Dujon B. 1986. Universal code equivalent of a yeast mitochondrial intron reading frame is expressed into *E. coli* as a specific double strand endonuclease. Cell 44:521–533.

Delport W, Poon AFY, Frost SDW, Kosakovsky Pond SL. 2010. Datamonkey 2010: a suite of phylogenetic analysis tools for evolutionary biology. Bioinformatics. 26:2455–2457.

Dieckmann C, Gandy B. 1987. Preferential recombination between GC clusters in yeast mitochondrial DNA. EMBO J. 6:4197–4203.

Dujon B, Slonimski PP, Weill L. 1974. Mitochondrial genetics IX: A model for recombination and segregation of mitochondrial genomes in *Saccharomyces cerevisiae*. Genetics. 78:415–437.

Dunn B, Richter C, Kvitek DJ, Pugh T, Sherlock G. 2012. Analysis of the *Saccharomyces cerevisiae* pan-genome reveals a pool of copy number variants distributed in diverse yeast strains from differing industrial environments. Genome Res. 22:908–924.

Edgar RC. 2004. MUSCLE: multiple sequence alignment with high accuracy and high throughput. Nucl Acids Res. 32:1792–1797.

Fay JC, Benavides JA. 2005. Evidence for domesticated and wild populations of *Saccharomyces cerevisiae*. PLoS Genet. 1:e5.

Fritsch ES, Chabbert CD, Klaus B, Steinmetz LM. 2014. A genome-wide map of mitochondrial DNA recombination in yeast. Genetics. 198:755–771.

Gallone B, Steensels J, Prahl T et al. 2016. Domestication and divergence of *Saccharomyces cerevisiae* beer yeasts. Cell. 166:1397–1410.

Gibson BR, Storgårds E, Krogerus K, Vidgren V. 2013. Comparative physiology and fermentation performance of Saaz and Frohberg lager yeast strains and the parental species *Saccharomyces eubayanus*. Yeast. 30:255–266.

Goddard MR, Burt A. 1999. Recurrent invasion and extinction of a selfish gene. Proc Natl Acad Sci U S A. 96:13880–13885.

Gonçalves M, Pontes A, Almeida P, Barbosa R, Serra M, Libkind D, Hutzler M, Gonçalves P, Sampaio JP. 2016. Distinct domestication trajectories in Top-Fermenting beer yeasts and wine yeasts. Curr Biol. Online.

González SS, Barrio E, Querol A. 2008. Molecular characterization of new natural hybrids between S. *cerevisiae* and S. *kudriavzevii* from brewing. Appl Environ Microbiol. 74:2314–2320.

Green DR, Reed JC. 1998. Mitochondria and Apoptosis. Science. 281:1309–1312.

Hao W, Richardson AO, Zheng Y, Palmer JD. 2010. Gorgeous mosaic of mitochondrial genes created by horizontal transfer and gene conversion. Proc Natl Acad Sci U S A. 107:21576–21581.

Hatefi Y. 1985. The mitochondrial electron transport and oxidative phosphorylation system. Annu Rev Biochem. 54:1015–1069.

Hill GE. 2015. Mitonuclear ecology. Mol Biol Evol. 32: 1917–1927.

Hittinger CT, Gonçalves P, Sampaio JP, Dover J, Johnston M, Rokas A. 2010. Remarkably ancient balanced polymorphisms in a multi-locus gene network. Nature. 464:54–58.

Hittinger CT. 2013. *Saccharomyces* diversity and evolution: a budding model genus. Trends in Genetics. 29:309–317.

Hou J, Friedrich A, Gounot JS, Schacherer J. 2015. Comprehensive survey of condition-specific reproductive isolation reveals genetic incompatibility in yeast. Nat Commun. 6:

Huson DH, Bryant D. 2006. Application of phylogenetic networks in evolutionary studies. Mol Biol Evol. 23:254–267.

Hyma KE, Fay JC. 2013. Mixing of vineyard and oak-tree ecotypes of *Saccharomyces cerevisiae* in North American vineyards. Mol Ecol. 22:2917–2930.

Kearse M, Moir R, Wilson A et al. 2012. Geneious Basic: An integrated and extendable desktop software platform for the organization and analysis of sequence data. Bioinformatics. 28:1647–1649.

Keeling P. 2009. Role of Horizontal Gene Transfer in the evolution of photosynthetic Eukaryotes and their plastids. In: Gogarten M, Gogarten J, Olendzenski L, editors. Horizontal Gene Transfer. Humana Press. p. 501–515.

Kosakovsky Pond SL, Posada D, Gravenor MB, Woelk CH, Frost SDW. 2006a. Automated phylogenetic detection of recombination using a genetic algorithm. Mol Biol Evol. 23:1891–1901.

Kosakovsky Pond SL, Posada D, Gravenor MB, Woelk CH, Frost SDW. 2006b. GARD: a genetic algorithm for recombination detection. Bioinformatics. 22:3096–3098.

Kück P, Meusemann K. 2010. FASconCAT, Version 1.0. Zool. Forschungsmuseum A. Koenig, Germany.

Kurtzman CP, Robnett CJ. 2003. Phylogenetic relationships among yeasts of the *‘Saccharomyces* complex’ determined from multigene sequence analyses. FEMS Yeast Res. 3:417–432.

Leducq JB, Charron G, Samani P et al. 2014. Local climatic adaptation in a widespread microorganism. Proc R Soc Lond B Biol Sci. 281:

Lee HY, Chou JY, Cheong L, Chang NH, Yang SY, Leu JY. 2008. Incompatibility of nuclear and mitochondrial genomes causes hybrid sterility between two yeast species. Cell. 135:1065–1073.

Legras JL, Merdinoglu D, Cornuet JM, Karst F. 2007. Bread, beer and wine: *Saccharomyces cerevisiae* diversity reflects human history. Mol Ecol. 16:2091–2102.

Librado P, Rozas J. 2009. DnaSP v5: a software for comprehensive analysis of DNA polymorphism data. Bioinformatics. 25:1451–1452.

Ling F, Shibata T. 2002. Recombination-dependent mtDNA partitioning: in vivo role of Mhr1p to promote pairing of homologous DNA. EMBO J. 21:4730–4740.

Ling F, Hori A, Shibata T. 2007. DNA recombination-initiation plays a role in the extremely biased inheritance of yeast [rho-] mitochondrial DNA that contains the replication origin ori5. Mol Cell Biol. 27:1133–1145.

Ling F, Mikawa T, Shibata T. 2011. Enlightenment of yeast mitochondrial homoplasmy: diversified roles of gene conversion. Genes. 2:169–190.

Ling F, Hori A, Yoshitani A, Niu R, Yoshida M, Shibata T. 2013. *Din7* and *Mhr1* expression levels regulate double-strand-break-induced replication and recombination of mtDNA at ori5 in yeast. Nucl Acids Res. 41:5799–5816.

Liti G, Peruffo A, James SA, Roberts IN, Louis EJ. 2005. Inferences of evolutionary relationships from a population survey of LTR-retrotransposons and telomeric-associated sequences in the *Saccharomyces sensu stricto* complex. Yeast. 22:177–192.

Liti G, Carter DM, Moses AM et al. 2009. Population genomics of domestic and wild yeasts. Nature. 458:337–341.

Lockshon D, Zweifel SG, Freeman-Cook LL, Lorimer HE, Brewer BJ, Fangman WL. 1995. A role for recombination junctions in the segregation of mitochondrial DNA in yeast. Cell. 81:947–955.

MacAlpine DM, Perlman PS, Butow RA. 1998. The high mobility group protein Abf2p influences the level of yeast mitochondrial DNA recombination intermediates in vivo. Proc Natl Acad Sci U S A. 95:6739–6743.

Martin DP, Lemey P, Lott M, Moulton V, Posada D, Lefeuvre P. 2010. RDP3: a flexible and fast computer program for analyzing recombination. Bioinformatics. 26:2462–2463.

Muller LAH, McCusker JH. 2009. A multispecies-based taxonomic microarray reveals interspecies hybridization and introgression in *Saccharomyces cerevisiae*. FEMS Yeast Res. 9:143–152.

Nakagawa K, Morishima N, Shibata T. 1992. An endonuclease with multiple cutting sites, Endo.SceI, initiates genetic recombination at its cutting site in yeast mitochondria. EMBO J. 11:2707–2715.

Nakhleh L. 2011. Evolutionary Phylogenetic Networks: Models and Issues. In: Heath SL, Ramakrishnan N, editors. Problem Solving Handbook in Computational Biology and Bioinformatics. Boston, MA: Springer US. p. 125–158.

Nunnari J, Marshall WF, Straight A, Murray A, Sedat JW, Walter P. 1997. Mitochondrial transmission during mating in *Saccharomyces cerevisiae* is determined by mitochondrial fusion and fission and the intramitochondrial segregation of mitochondrial DNA. Mol Biol Cell. 8:1233–1242.

Okuno M, Kajitani R, Ryusui R, Morimoto H, Kodama Y, Itoh T. 2016. Next-generation sequencing analysis of lager brewing yeast strains reveals the evolutionary history of interspecies hybridization. DNA Res. 23:67–80.

Paliwal S, Fiumera AC, Fiumera HL. 2014. Mitochondrial-nuclear epistasis contributes to phenotypic variation and coadaptation in natural isolates of *Saccharomyces cerevisiae*. Genetics. 198:1251–1265.

Pérez-Través L, Lopes CA, Querol A, Barrio E. 2014. On the complexity of the *Saccharomyces bayanus* taxon: hybridization and potential hybrid speciation. PLoS ONE. 9:e93729.

Peris D, Belloch C, Lopandic K, Álvarez-Pérez JM, Querol A, Barrio E. 2012a. The molecular characterization of new types of S. *cerevisiae* × S. *kudriavzevii* hybrid yeasts unveils a high genetic diversity. Yeast. 29:81–91.

Peris D, Lopes CA, Arias A, Barrio E. 2012b. Reconstruction of the evolutionary history of *Saccharomyces cerevisiae* × S. *kudriavzevii* hybrids based on multilocus sequence analysis. PLoS ONE. 7:e45527.

Peris D, Lopes CA, Belloch C, Querol A, Barrio E. 2012c. Comparative genomics among *Saccharomyces cerevisiae* × *Saccharomyces kudriavzevii* natural hybrid strains isolated from wine and beer reveals different origins. BMC Genomics. 13:407.

Peris D, Sylvester K, Libkind D, Gonçalves P, Sampaio JP, Alexander WG, Hittinger CT. 2014. Population structure and reticulate evolution of *Saccharomyces eubayanus* and its lager-brewing hybrids. Mol Ecol. 23:2031–2045.

Peris D, Langdon Q, Moriarty R et al. 2016. Complex ancestries of lager-brewing hybrids were shaped by standing variation in wild yeast *Saccharomyces eubayanus*. Plos Genetics 12 (7): e1006155

Posada D. 2008. jModelTest: phylogenetic model averaging. Mol Biol Evol. 25:1253–1256.

Procházka E, Franko F, Poláková S, Sulo P. 2012. A complete sequence of *Saccharomyces paradoxus* mitochondrial genome that restores the respiration in *S. cerevisiae*. FEMS Yeast Res. 12:819–839.

Querol A, Barrio E, Huerta T, Ramon D. 1992. Molecular monitoring of wine fermentations conducted by active dry yeast strains. Appl Environ Microbiol. 58:2948–2953.

Rodríguez ME, Pérez-Través L, Sangorrín MP, Barrio E, Lopes CA. 2014. *Saccharomyces eubayanus* and *Saccharomyces uvarum* associated with the fermentation of *Araucaria araucana* seeds in Patagonia. FEMS Yeast Res. 14:948–965.

Sampaio JP, Gonçalves P. 2008. Natural populations of *Saccharomyces kudriavzevii* in Portugal are associated with oak bark and are sympatric with S. *cerevisiae* and S. *paradoxus*. Appl Environ Microbiol. 74:2144–2152.

Scannell DR, Zill OA, Rokas A, Payen C, Dunham MJ, Eisen MB, Rine J, Johnston M, Hittinger CT. 2011. The awesome power of yeast evolutionary genetics: new genome sequences and strain resources for the *Saccharomyces sensu stricto* genus. G3 1:11–25.

Schacherer J, Shapiro JA, Ruderfer DM, Kruglyak L. 2009. Comprehensive polymorphism survey elucidates population structure of *Saccharomyces cerevisiae*. Nature. 458:342–345.

Schmidt HA, Strimmer K, Vingron M, von Haeseler A. 2002. TREE-PUZZLE: maximum likelihood phylogenetic analysis using quartets and parallel computing. Bioinformatics. 18:502–504.

Séraphin B, Simon M, Faye G. 1987. The mitochondrial reading frame *RF3* is a functional gene in *Saccharomyces uvarum*. J Biol Chem. 262:10146–10153.

Shimodaira H, Hasegawa M. 1999. Multiple comparisons of Log-Likelihoods with applications to Phylogenetic inference. Mol Biol Evol. 16:1114–1116.

Sickmann A, Reinders J, Wagner Y et al. 2003. The proteome of *Saccharomyces cerevisiae* mitochondria. Proc Natl Acad Sci U S A. 100:13207–13212.

Stamatakis A. 2014. RAxML version 8: a tool for phylogenetic analysis and post-analysis of large phylogenies. Bioinformatics. 30:1312–1313.

Starkov AA. 2008. The role of mitochondria in reactive oxygen species metabolism and signaling. Annals of the New York Academy of Sciences. 1147:37–52.

Strathern JN, Jones EW, Broach J.R., Dujon B. 1981. Mitochondrial genetics and functions. In: Strathern JN, Jones EW, Broach JR, editors. Molecular Biology of the Yeast Saccharomyces Life Cycle and Inheritance. Cold Spring Harbor, NY: Cold Spring Harbor Laboratory Press. p. 505–635.

Strimmer K, Rambaut A. 2002. Inferring confidence sets of possibly misspecified gene trees. Proceedings Biological sciences / The Royal Society. 269:137–142.

Strope PK, Skelly DA, Kozmin SG, Mahadevan G, Stone EA, Magwene PM, Dietrich FS, McCusker JH. 2015. The 100-genomes strains, an S. *cerevisiae* resource that illuminates its natural phenotypic and genotypic variation and emergence as an opportunistic pathogen. Genome Res. 25:1–13.

Sun S, Evans BJ, Golding GB. 2011. “Patchy-Tachy” leads to false positives for recombination. Mol Biol Evol. 28:2549–2559.

Sylvester K, Wang QM, James B, Mendez R, Hulfachor AB, Hittinger CT. 2015. Temperature and host preferences drive the diversification of *Saccharomyces* and other yeasts: a survey and the discovery of eight new yeast species. FEMS Yeast Res. 15:1–16.

Tamura K, Peterson D, Peterson N, Stecher G, Nei M, Kumar S. 2011. MEGA5: Molecular Evolutionary Genetics Analysis using Maximum Likelihood, evolutionary distance, and Maximum Parsimony methods. Mol Biol Evol. 28:2731–2739.

Taylor JW. 1986. Fungal evolutionary biology and mitochondrial DNA. Exp Mycol. 10:259–269.

Waterhouse AM, Procter JB, Martin DMA, Clamp M, Barton GJ. 2009. Jalview Version 2 -a multiple sequence alignment editor and analysis workbench. Bioinformatics. 25:1189–1191.

Wolters JF, Chiu K, Fiumera HL. 2015. Population structure of mitochondrial genomes in *Saccharomyces cerevisiae*. BMC Genomics. 16:451.

Wu B, Buljic A, Hao W. 2015. Extensive horizontal transfer and homologous recombination generate highly chimeric mitochondrial genomes in yeast. Mol Biol Evol. 32:2559–2570.

Wu B, Hao W. 2015. A dynamic mobile DNA family in the yeast mitochondrial genome. G3: Genes|Genomes|Genetics. 5:1273–1282.

Yang J, Mohr G, Perlman PS, Lambowitz AM. 1998. Group II intron mobility in yeast mitochondria: target DNA-primed reverse transcription activity of ai1 and reverse splicing into DNA transposition sites in vitro1. J Mol Biol. 282:505–523.

Zeyl C, Andreson B, Weninck E. 2005. Nuclear-mitochondrial epistasis for fitness in *Saccharomyces cerevisiae*. Evolution. 59:910–914.

